# Scaffold pore geometry guides gene regulation and bone-like tissue formation in dynamic cultures

**DOI:** 10.1101/2020.04.24.060525

**Authors:** Marina Rubert, Jolanda Rita Vetsch, Iina Lehtoviita, Marianne Sommer, Feihu Zhao, André R Studart, Ralph Müller, Sandra Hofmann

## Abstract

Cells sense and respond to scaffold pore geometry and mechanical stimuli. Many fabrication methods used in bone tissue engineering render structures with poorly controlled pore geometries. Given that cell-scaffold interactions are complex, drawing a conclusion on how cells sense and respond to uncontrolled scaffold features under mechanical loading is difficult. Here, monodisperse templated scaffolds (MTSC) were fabricated and used as a well-defined porous scaffolds to study the effect of dynamic culture conditions on bone-like tissue formation. Human bone marrow derived stromal cells were cultured on MTSC or conventional salt-leached scaffolds (SLSC) for up to 7 weeks, either under static or dynamic conditions (wall shear stress (WSS) using spinner flask bioreactors). The influence of controlled spherical pore geometry of MTSC subjected to static or dynamic conditions on osteoblast cells differentiation, bone-like tissue formation, structure and distribution was investigated. WSS generated within the two idealized geometrical scaffold features was assessed. Distinct response to fluid flow in osteoblast cell differentiation were shown to be dependent on scaffold pore geometry. As revealed by collagen staining and micro-computed tomography images, dynamic conditions promoted a more regular extracellular matrix (ECM) formation and mineral distribution in both scaffold types compared to static conditions. The results showed that regulation of bone-related genes and the amount and the structure of mineralized ECM were dependent on scaffold pore geometry and the mechanical cues provided by the two different culture conditions. Under dynamic conditions, SLSC favored osteoblast cell differentiation and ECM formation, while MTSC enhanced ECM mineralization. The spherical pore shape in MTSC supported a more trabecular bone-like structure under dynamic conditions compared to MTSC statically cultured or to SLSC under either static or dynamic conditions. These results suggest that cell activity and bone-like tissue formation is driven not only by the pore geometry but also by the mechanical environment. This should be taken into account in the future design of complex scaffolds, which should favor cell differentiation while guiding the formation, structure and distribution of the engineered bone tissue. This could help to mimic the anatomical complexity of the bone tissue structure and to adapt to each bone defect needs.

**Impact statement:** Aging of the human population leads to an increasing need for medical implants with high success rate. We provide evidence that cell activity and the amount and structure of bone-like tissue formation is dependent on the scaffold pore geometry and on the mechanical environment. Fabrication of complex scaffolds comprising concave and planar pore geometries might represent a promising direction towards the tunability and mimicry the structural complexity of the bone tissue. Moreover, the use of fabrication methods that allow a systematic fabrication of reproducible and geometrically controlled structures would simplify scaffold design optimization.

## 1. Introduction

Restoring large bone defects remains a clinical challenge [1]. Although autologous bone grafts are the gold standard treatment, problems related to donor site morbidity and limited amount of available bone tissue remain a major drawback [2]. Implantation of biocompatible scaffolds that support the migration of the patients’ own cells, their proliferation and new bone tissue formation (BTF) at the defect site could represent a cost- and time-effective alternative [3].

Numerous *in vitro* and *in vivo* studies have focused on the optimization of scaffold architectures to improve the biological response and to fulfill a variety of functional requirements [4–10]. Despite evidence that scaffold architecture affects its performance and cellular response, it is still not clear what the best pore size, shape and interconnectivity of the scaffold architecture should be. One reason is that the selection of an optimal bone graft varies from patient to patient and relies on, the defect size and shape, on the anatomical location and the specific biomechanical or functional requirements at that site [11]. Given that cell-scaffold interactions are complex and that the biological mechanism by which the scaffold mediates BTF is still missing [12], drawing a general conclusion on how cells sense and respond to a certain architectural scaffold feature is difficult [11]. This becomes even more challenging if one considers the poorly controlled structure of scaffolds typically used for bone tissue engineering. Many of the fabrication methods used in bone tissue engineering (e.g. salt leaching, gas foaming, lyophilization) often fail to allow an independent variation of individual geometric parameters [13]. To better understand cell-scaffold interactions and thus advance on the development of new strategies to enhance BTF, it is necessary to use methods that enable the fabrication of reproducible and geometrically controlled structures. Moreover, the use of fabrication methods that allow a systematic fabrication and evaluation of the influence of individual scaffold parameters on cell behavior and promoting BTF would simplify scaffold optimization to fulfill the requirements of the patient and the mechanical conditions at the defect site.

The next generation of three dimensional (3D) scaffolds to enhance *in vivo* BTF aims at supporting the amount of bone tissue formed but also at closely mimicking the anatomical organization of bone tissue and its extracellular matrix (ECM) at the defect site [14]. Attention has been given to the stimulating and guiding role of the geometrical design of scaffolds [15–25]. *In vitro* and *in vivo* studies have suggested that the structure of engineered bone-like tissue can be modulated by controlling the scaffold pore diameter [21, 24, 26, 27], the channel orientation [16] and the porous scaffold architecture [17, 20]. *In vitro,* two-dimensional pore geometry and surface curvature have further shown to have a strong influence on cell viability [18], cell proliferation [15, 22, 25] and differentiation [19, 25] or overall tissue formation [15, 22, 25]. Cell-independent mineral nucleation was shown to be regulated by material surface curvature [19, 28]. It has been suggested that controlling the 3D scaffold pore geometry has a major impact on the bioactivity of a scaffold towards stem cell differentiation into bone tissue forming cells and subsequent ECM formation and mineralization [23]. In that study, silk fibroin (SF), a widely-used material for bone tissue engineering applications [29–31], was selected as material for scaffold fabrication. A new fabrication method for SF scaffolds with a highly controlled and monodisperse porous architecture was developed [23], which were referred to by the authors as inverse opal scaffolds. Here we name them, for consistency to the salt leached scaffolds based on the porogen, monodisperse template scaffolds (MTSC). A systematic evaluation of physical features and biological response of MTSC was performed relative to conventional salt-leached SF scaffolds (SLSC). The inverse opal method used poly(caprolactone) (PCL) as spherical monodisperse pore template. MTSC result in a close packed and regular spherical pore arrangement. While keeping comparable stiffness, roughness and mean pore diameter, MTSC showed a monodisperse pore diameter distribution, regular spherical shapes and lower overall porosity compared to SLSC [23]. While no differences were observed in cell proliferation, human bone marrow derived stromal cells (hBMSCs) cultured on MTSC under static conditions produced significantly more matrix mineralization compared to cells cultured on SLSC [23].

In addition to scaffold stiffness and topography (pore geometry), cells sense and respond to changes in their mechanical environment [32]. Spinner flask bioreactors have been used in tissue engineering to overcome nutrient constraints through large scaffolds but also to induce shear stresses *in vitro* through generation of increased fluid flow. Fluid-induced shear stress has been shown to alter the cytoskeletal composition of osteoblastic cells [33] and to induce osteoblast cell differentiation [34] and ECM mineralization [35–37]. Although substrate geometry and shear stresses have been shown to guide the osteoblast cell differentiation, so far studies were limited to the investigation of these two components independently.

Our goal was to assess the influence different scaffold pore geometries on fluid flow and bone-like tissue formation. More specifically, we investigated whether dynamic cell culture conditions, generated by culturing hBMSC in spinner flask bioreactors, play a role on the osteogenic cell fate and on the spatial organization of mineralized tissue within 3D SF scaffolds exhibiting either spherical and monodisperse (MTSC) or faceted and polydisperse (SLSC) porous architectures [23]. Furthermore, we investigated whether scaffold type and shear stress could have a synergistic effect on increasing cell differentiation and mineralization.

## 2. Material and methods

### 2.1. Preparation of SF solution

SF solution was prepared as previously described [38]. Briefly, silk cocoons from *B. mori* (Trudel Inc.) were boiled twice in 0.02 M Na_2_CO_3_ solution for 1 hour and then washed with ultrapure water (UPW). Purified SF was solubilized in 9 M lithium bromide solution (ThermoScientific) and dialyzed (3.5K molecular weight cut-off, ThermoScientific) against UPW for 36 hours. After freezing at −80°C, the frozen SF solution was freeze-dried for 5 days. For SLSC, lyophilized SF was dissolved at 17% (w/v) in 1,1,1,3,3,3-hexafluoroisopropanol (HFIP) (FluoroChem Ltd.). For MTSC, lyophilized SF was dissolved at 12% (w/v) in HFIP.

### 2.2. Production of SLSC

Sodium chloride (NaCl) crystals with a granule diameter between 300-400 μm were used as a faceted polydisperse porogen. 1 ml of the 17% (w/v) SF-HFIP solution was added to 2.5 g of NaCl crystals. The HFIP was allowed to evaporate for 3 days. β-sheet conformation was induced by immersing the SF-salt blocks into 90% (v/v) methanol (Merck) in UPW for 30 minutes and blocks were subsequently dried overnight [39]. NaCl was leached by washing the SF-salt blocks in UPW for 2 days. SF blocks were cut and punched (Kai medical) into disks (3 mm height, 5 mm diameter). SLSC were sterilized in phosphate buffered saline (PBS; Medicago AB) at 121°C, 1 bar for 20 minutes. The SLSC’s fabrication process is depicted in figure 1A.

**Fig. 1.**
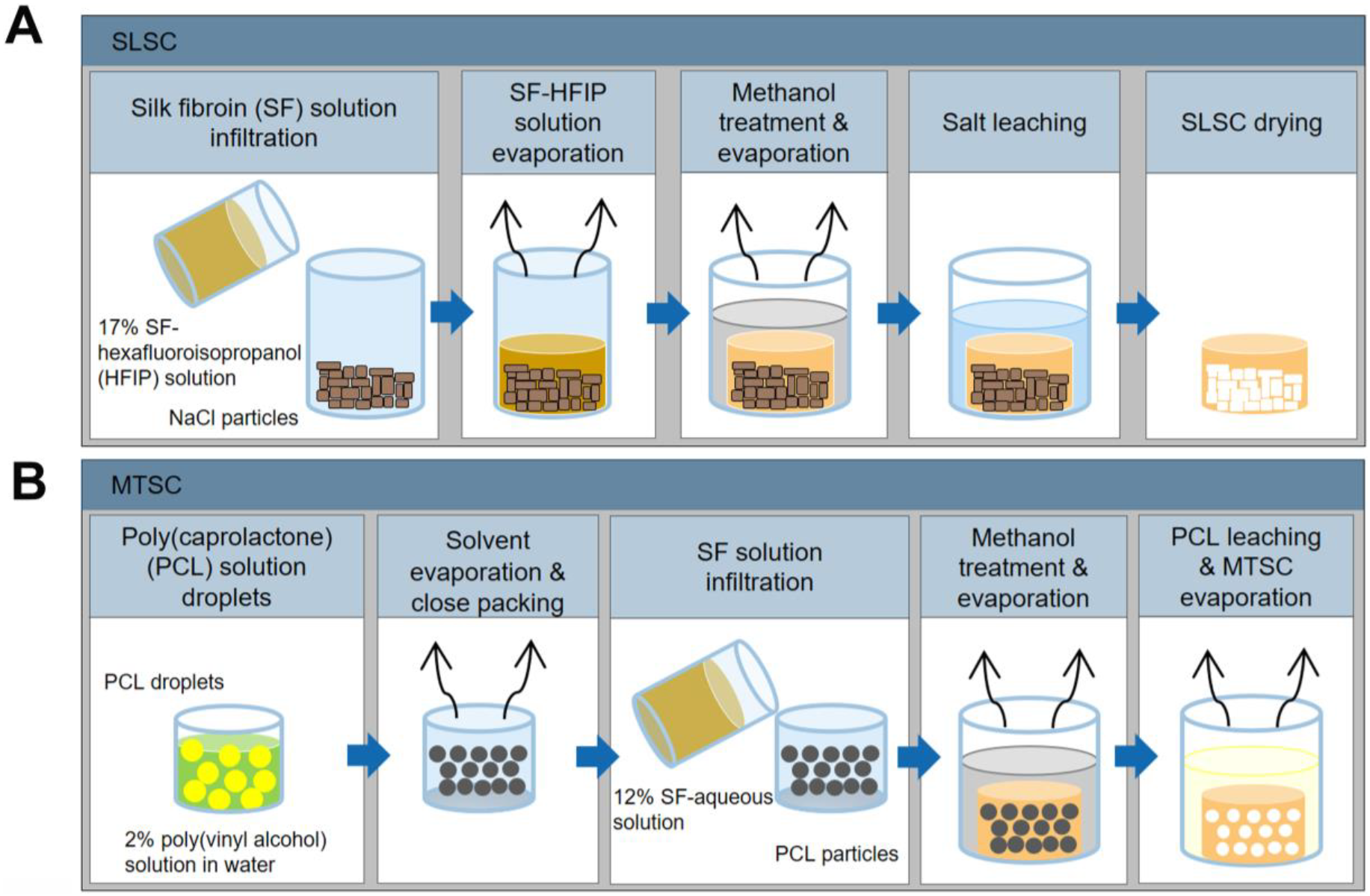
Schematic illustration of the production process of salt-leached scaffolds (SLSC) (A) and inverse opal scaffolds (MTSC) (B).

### 2.3. Production of MTSC

A 10% (w/w) PCL in dichloromethane solution (SIGMA) was emulsified in a 2% (w/w) poly(vinyl alcohol) (PVA; SIGMA) aqueous solution within a glass capillary microfluidic device to produce monodisperse droplets [23]. The inner and the outer flow rates were adjusted to 5 ml/hr and 8 ml/hr, respectively. The droplets were collected in an excess of PVA solution and left resting for approximately five weeks until dichloromethane was completely evaporated. The particles were washed 5 times in water and once in ethanol. Subsequently, particles were placed in moulds (7 mm diameter, 3 mm height) and slightly agitated to close pack and dry. The particles were then heated to 57°C at 25% relative humidity for 1 hour to form necks between the particles. 12% SF solution was infiltrated into each particle pack and let dry. β-sheet conformation was induced by immersing the SF-salt blocks into 90% (v/v) methanol in UPW for 1 hour [40]. Following 8 hours in dichloromethane to remove the porogen particles, samples were washed once in ethanol to remove any solvent residues and left to dry. Finally, MTSC were sterilized in PBS at 121°C, 1 bar for 20 min. Due to the scaffold fragility after autoclaving, scaffolds were not punched further out. The MTSC’s fabrication process is depicted in figure 1B.

### 2.4. Computational fluid dynamics (CFD) simulation

To quantify the wall shear stress (WSS) generated within the two scaffold types, a CFD model was developed based on the spinner flask bioreactor (figure 2A). For saving computational cost, a computer-aided design (CAD) was used for creating idealized scaffold geometries (figures 2B and C). Based on the size and shape of the NaCl and PCL particles used as porogen in SLSC and MTSC, the scaffold geometric features for CAD were determined as: a pore diameter (*d*) of 330 μm for both scaffold types; a cubical pore shape for SLSCs and a spherical pore shape for MTSCs. According to previous measurements, the average porosity (*φ*) of MTSC and SLSC was 84.7% and 92.8%, respectively [23]. Two CFD domains were meshed with 1,187,159 tetrahedron elements (SLSC scaffold) and 2,650,122 tetrahedron elements (MTSC scaffold), respectively, using the same mesh strategy as previously [41]. The density (*ρ*) and dynamic viscosity (*μ*) of the culture medium were 1000 kg/m^3^ and 1.45 mPa·s, respectively (Olivares et al., 2009). The applied shear stress transport (SST) turbulence model was the same as described in Melke et al. [41]. The boundary conditions are illustrated in figure 2A. To solve the CFD model, commercial finite volume method codes (CFX, ANSYS Inc., USA) were used. The model was solved under steady state when reaching the convergence criteria: the root-mean-square of the residuals of the mass and the momentum < 10^−4^.

**Fig. 2.**
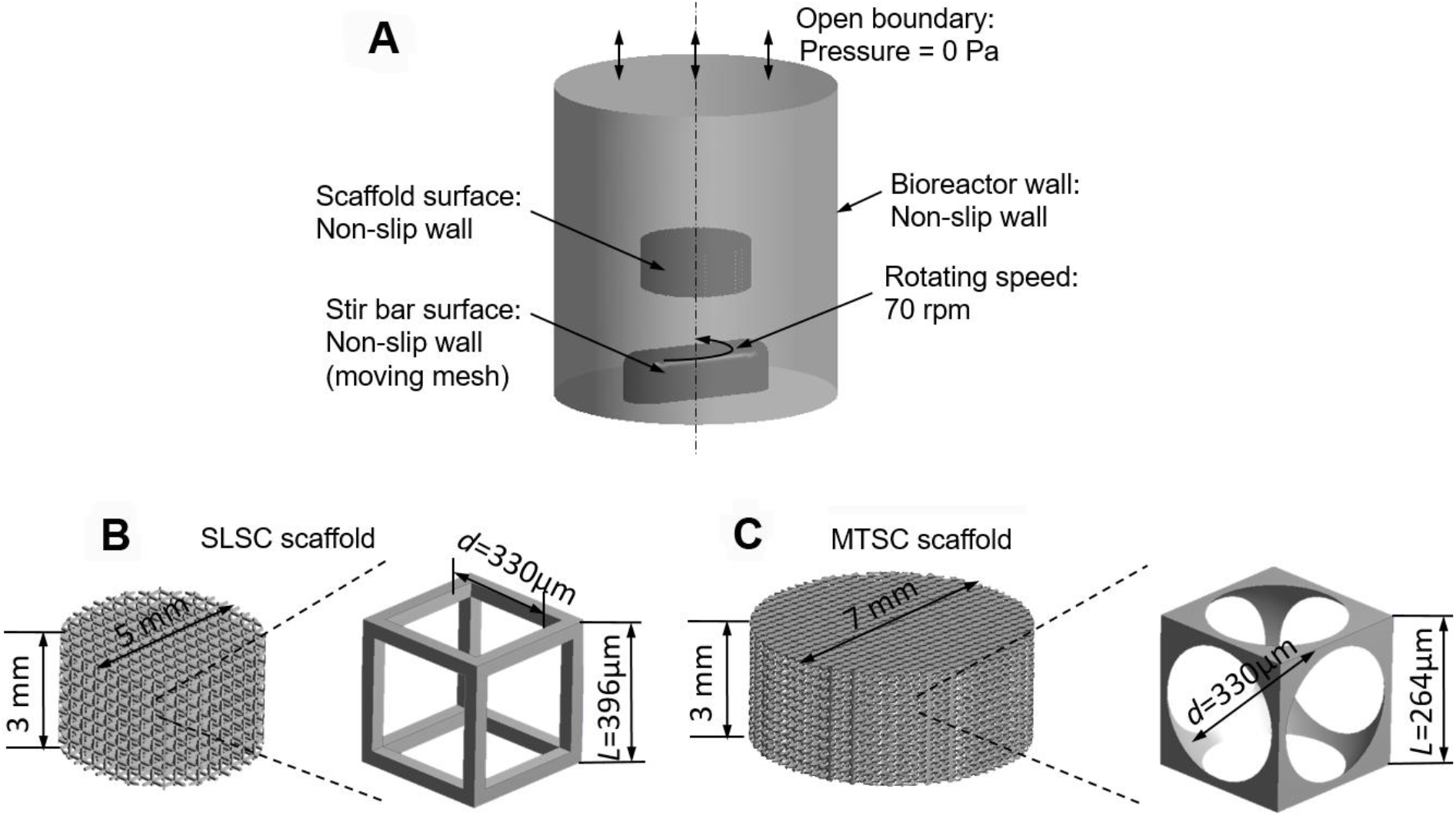
(A) Fluid domain of CFD model and illustration of boundary conditions; CAD geometries of (B) SLSC scaffold that is assembled by repeating units with cubical pores (pore size *d* = 330 μm, porosity *φ* = 92.8%); (C) MTSC scaffold that is assembled by repeating units with spherical pores (pore size *d* = 330 μm, porosity *φ* = 84.7%).

### 2.5. Cell culture

hBMSCs were isolated and characterized from bone marrow (Lonza) as previously described [20]. hBMSCs (passage 3) were expanded at 37°C and 5% CO_2_ in expansion medium (Dulbecco’s modified Eagle’s medium (DMEM; GIBCO) containing 10% fetal bovine serum (FBS; GIBCO), 1% Penicillin/Streptomycin/Fungizone (P/S/F; Invitrogen), 1% non-essential amino acids (Invitrogen) and 1 ng/ml basic fibroblast growth factor (Invitrogen)). After one week, 1*10^6^ cells dispersed in 20 μl medium were seeded on each scaffold. After cell incubation for 90 min under standard cell culture conditions (37°C, 5% CO_2_), the scaffolds were transferred into static (static conditions) or spinner flask bioreactors (dynamic conditions). Each bioreactor contained three scaffolds. Cells were cultured in 5 ml osteogenic media (DMEM, 10% FBS, 1% P/S/F, 50 μg/ml L-ascorbic acid (SIGMA), 100 nM dexamethasone (SIGMA), 10 mM beta-glycerolphosphate (SIGMA)) for 7 weeks either under static or dynamic culture conditions. The bioreactor caps were loosened to allow gas exchange. Culture media was changed every 2-3 days.

### 2.6. Dynamic culture conditions

Spinner flask bioreactors were chosen as a dynamic culture to induce fluid flow through and around the cell-scaffold construct. The culture medium was stirred through a magnet at approximately 70 rpm.

### 2.7. Histology

Constructs were fixed in 4 % formaldehyde overnight at room temperature. Samples were dehydrated, embedded in paraffin and cut into 10 μm thick sections. Weigert’s hematoxylin and Sirius red were used to stain the cell nuclei and collagen simultaneously. Images were taken using a Slide Scanner Pannoramic 250 (3DHISTECH). To improve the visualization of the histological staining, contrast and brightness of figure 5 and 6 were adapted using the same setting parameters in all images. Figures show representative images per group.

### 2.8. RNA isolation and Real-time PCR analysis

Total RNA was isolated using Trizol reagent (Invitrogen), according to the manufacturer’s protocol. Each scaffold was disintegrated at 25’000 rpm for 10 seconds per cycle (6 cycles) using steel balls and a Minibead Beater (Biospec). Samples were placed on ice in between cycles. Total RNA (1 μg) was reverse transcribed to cDNA at 42°C for 60 min using the High Capacity RNA-to-cDNA kit (Applied Biosystems), according to the manufacture’s protocol. 10 μl of cDNA was diluted into 10 μl of RNAse free water (SIGMA).

Real-time PCR (Biorad CFX96) was performed using TaqMan probe detection (Applied Biosystems). Real-time PCR was done for glyceraldehyde-3-phosphate dehydrogenase (GAPDH, Hs02758991_g1), collagen type I (COL1A2-I, Hs01028956_m1), alkaline phosphatase (ALPL, Hs01029144_m1) and osteocalcin (BGLAP, Hs01587814_g1). A negative control without cDNA template was run for each assay. Values were normalized to GAPDH mRNA levels as an internal control and were presented as fold change compared to SLSC in static conditions. Data was analyzed using the comparative Cq method (2-ΔCCq) [42].

### 2.9. Micro-computed tomography

For quantification and visualization of the engineered mineralized bone-like tissue, scaffolds were analyzed using a micro-CT imaging system (μCT40, Scanco Medical AG). Samples were non-invasively scanned weekly for up to 7 weeks. All bioreactors were scanned at 36 μm resolution, 45 kVp and 177μA. An integration time of 200 ms and 2-fold data averaging was applied. Cylindrical volumes of interest were selected for the analysis. A constraint Gaussian filter of 1.2 and a filter support of 1 was used to suppress noise. Mineralized tissue was segmented from non-mineralized tissue using a global thresholding procedure [43]. All samples were thresholded at a grey value of 149 (corresponding to a density of 89.1 mg HA per cm^3^). Quantitative histomorphometry was performed on all scaffolds to assess engineered bone-like tissue volume fraction (BV/TV), bone-like tissue surface-to-bone volume (BS/BV), trabecular thickness (Tb.Th.), trabecular separation (Tb.Sp.) and trabecular number (Tb.N.) [44, 45].

### 2.10. Statistics

Data (n=3) represent the mean ± standard deviation (SD). SPSS® 22.0 was used for statistical analysis. A one-way ANOVA with post-hoc Bonferroni correction was performed to compare differences among groups. P-values ≤ 0.05 were considered statistically significant. Figure 5 shows upper median samples based on the BV/TV values of the whole scaffold.

## 3. Results

### 3.1. CFD simulation

The CFD simulation shows the fluid velocity gradient within the two scaffold types in a spinner flask bioreactor environment (figure 3). For both, the bottom and peripheral regions of the scaffolds were subjected to a higher fluid velocity compared to the center region. This resulted in higher WSS in the bottom and peripheral scaffold regions than in the central scaffold regions for both scaffold types (figure 4). Comparing the WSS between the two scaffold types, the SLSC scaffold had a higher average WSS (0.81 mPa) than MTSC scaffold (0.42 mPa). For the SLSC, 59.71% and 38.50% of the scaffold surface area were subjected to a WSS in the range of 0.11 – 0.55 mPa and 0.55 – 10 mPa, respectively. For the MTSC, a surface area fraction of 70.00% and 12.43% in the WSS range of 0.11 – 0.55 mPa and 0.55 – 10 mPa, respectively. With the previously reported WSS thresholds for osteogenesis (0.11-10 mPa) [46] and mineralization of ECM (> 0.55 mPa) [47], this would predict that cells attached to approximately 60% (SLSC) strut surface and 70.00% (MTSC) strut surface area would be stimulated to differentiate towards the osteogenic lineage but without (yet) showing mineralization. This would happen in the upper centre region for SLSC and throughout the whole volume except the very bottom of the scaffold for the MTSC (light blue regions in figure 4). The model also predicted mineralized matrix formation in the bottom and peripheral regions for both pore geometries (purple regions in figure 4).

**Fig. 3.**
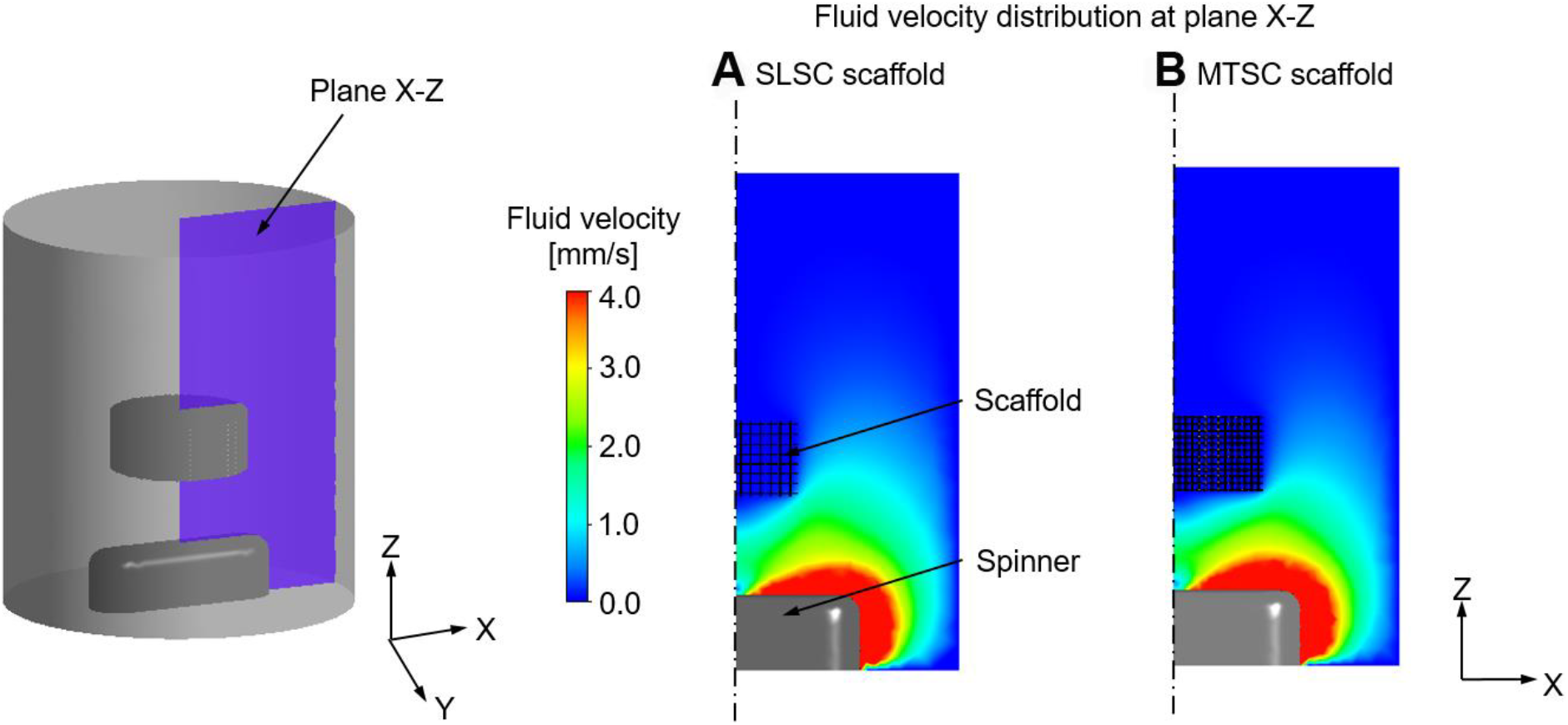
Cross-sectional view of the fluid velocity distribution within the spinner flask for (A) SLSC scaffold and (B) MTSC scaffold.

**Fig. 4.**
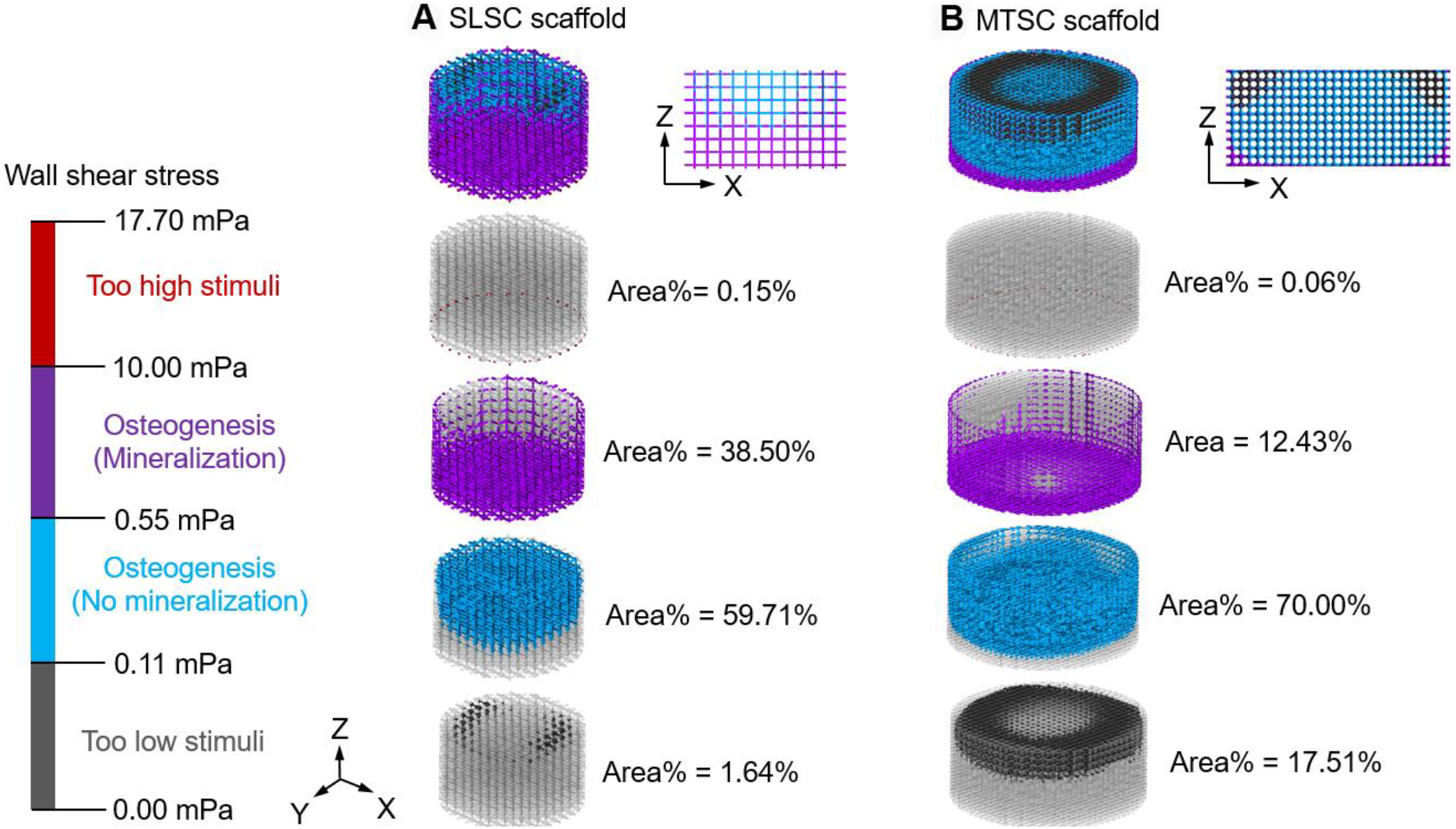
Prediction of regions within (A) SLSC and (B) MTSC scaffolds receiving different mechanical stimuli within a spinner flask bioreactor.

### 3.2. Cell distribution, ECM formation

Hematoxylin staining revealed that, while cells could infiltrate and distribute regularly in SLSC both under static and dynamic conditions (figure 5A, B), homogenous cell distribution in MTSC was only observed in the dynamic culture (figure 5D). In addition, cells migrated away from the longitudinal axis of the pore in MTSC (figure 5C-D, figure S2). Sirius red staining showed the formation of a collagenous ECM that was more evenly distributed in SLSC compared to MTSC, both under static and dynamic conditions. In line with the computational simulation of the WSS (figure 4), both scaffold pore geometries showed a denser collagenous ECM formation resulting from stimulated osteogenic differentiation in the upper part of the scaffold (figure 6). Acellular SLSC and MTSC were stained in parallel to count for potential false positive staining (data not shown).

**Fig. 5.**
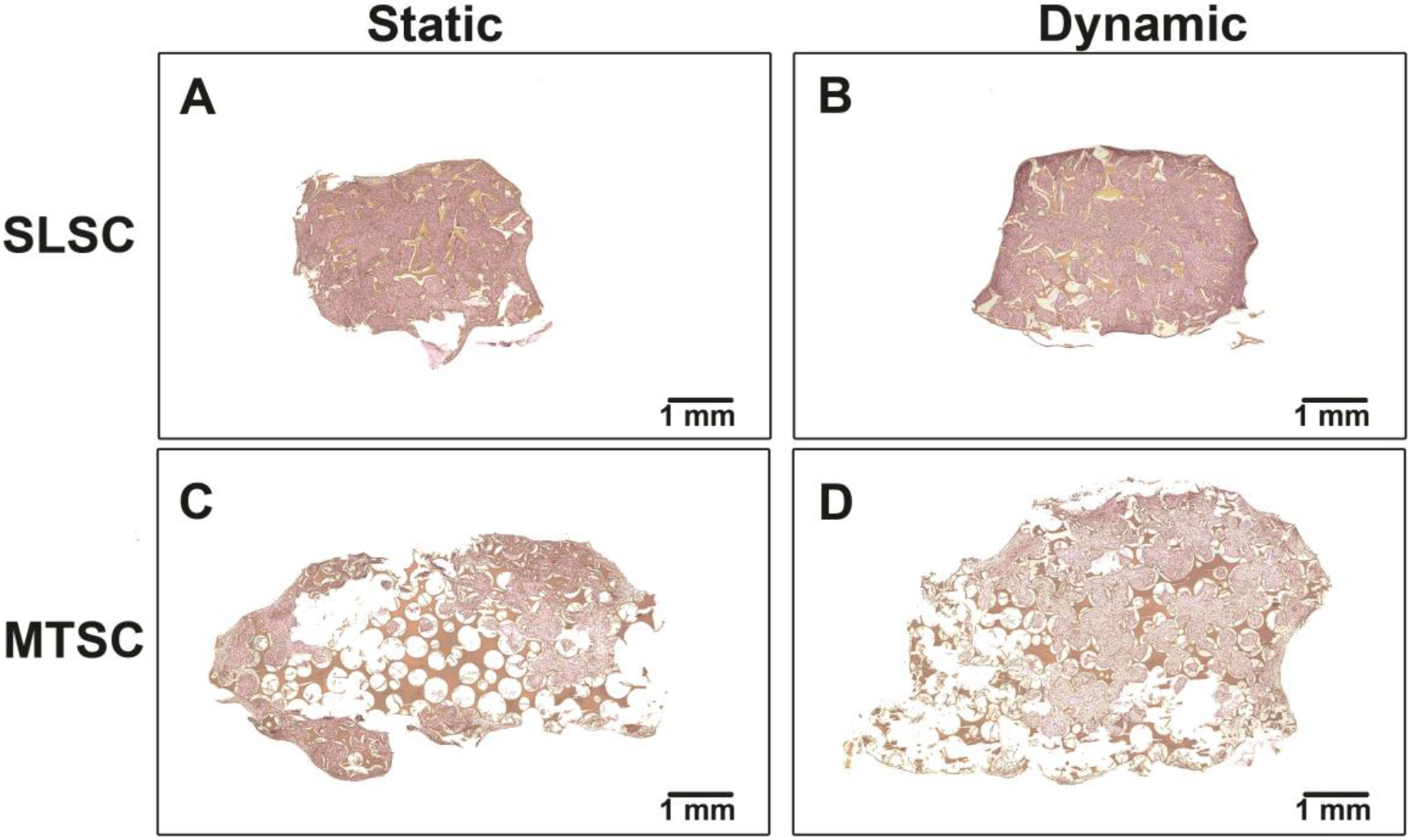
Overview of the cell distribution and formation of a collagenous ECM along the scaffold size. Weigert’s haematoxylin staining (cell nuclei - dark purple spots) and Sirius red staining (collagenous ECM - pink) on hBMSCs cultured on SLSC (A, B) and MTSC (C, D) either under static (A, C) or dynamic conditions (B, D). Scaffolds are shown in orange color. Scale bar lengths are 1 mm.

**Fig. 6.**
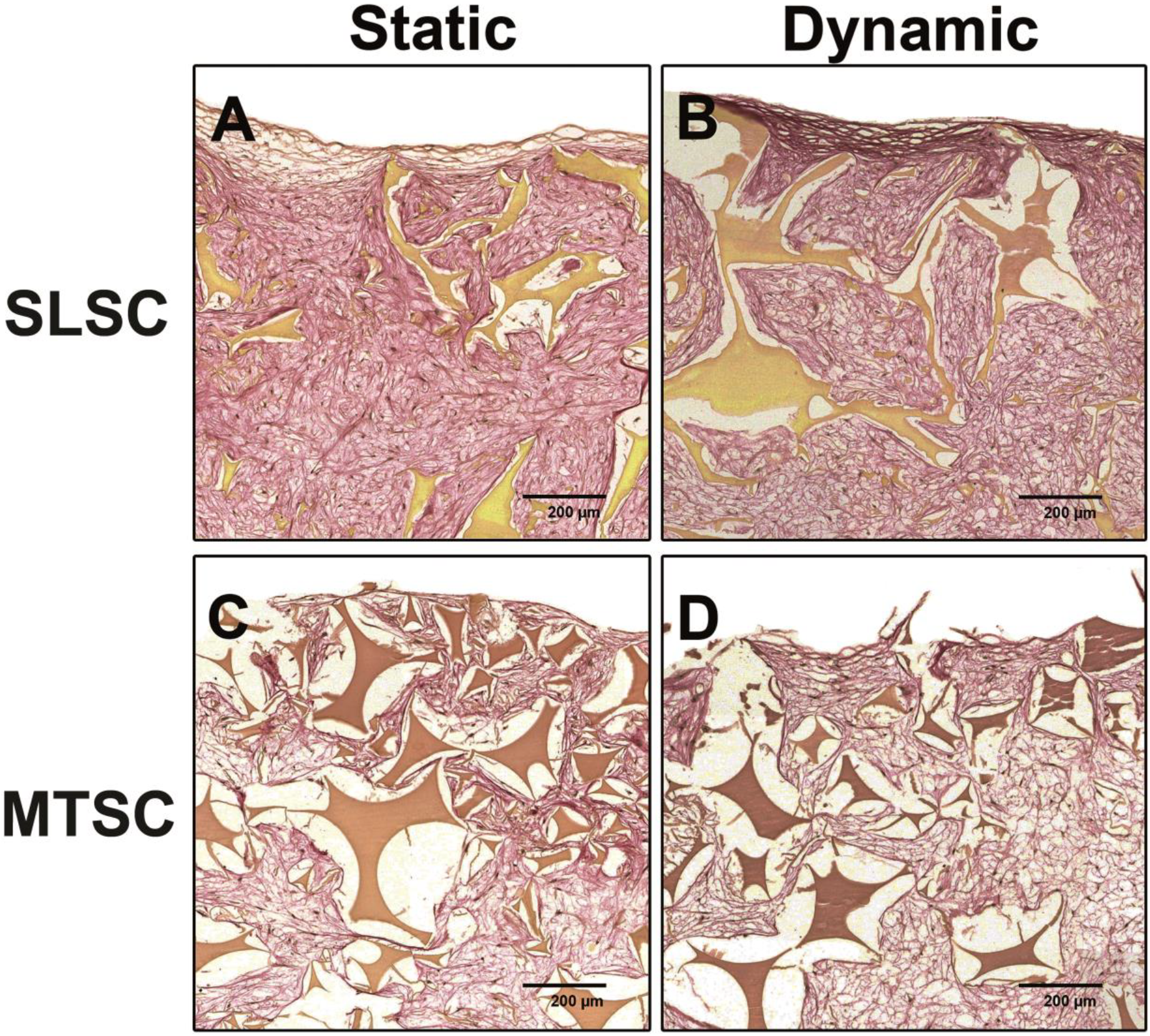
Magnified view of the formation of a collagenous ECM at the scaffold surface. Weigert’s haematoxylin staining (cell nuclei - dark purple spots) and Sirius red staining (collagenous ECM - pink) on hBMSCs cultured on SLSC (A, B) and MTSC (C, D) either under static (A, C) or dynamic conditions (B, D). Scaffolds are shown in orange color. Image is taken from the top-middle region of the scaffold. Scale bar lengths are 200 μm.

### 3.3. Bone-related mRNA expression levels

Dynamic conditions up-regulated the expression of osteoblastic differentiation markers in SLSC and MTSC compared to static conditions. This was evidenced by a significant increase of the ALPL mRNA levels by 3.1-fold (p = 0.00045, figure 7B) or of the BGLAP mRNA levels by 3.5-fold (p = 0.00500, figure 7C) in cells cultured on MTSC and on SLSC, respectively, compared to static conditions. The expression of these genes was also influenced by pore geometry. Under dynamic conditions, these genes were differentially expressed in both scaffolds. After 7 weeks of culture, ALPL mRNA levels were markedly up-regulated in MTSC compared to SLSC (8.4-fold, p = 0.00002) while BGLAP mRNA levels were significantly down-regulated (51-fold, p = 0.00043, figure 7B, 7C). Under static conditions, COL1A2 mRNA levels were up-regulated on SLSC compared to MTSC (p = 0.03, figure 7A).

**Fig. 7.**
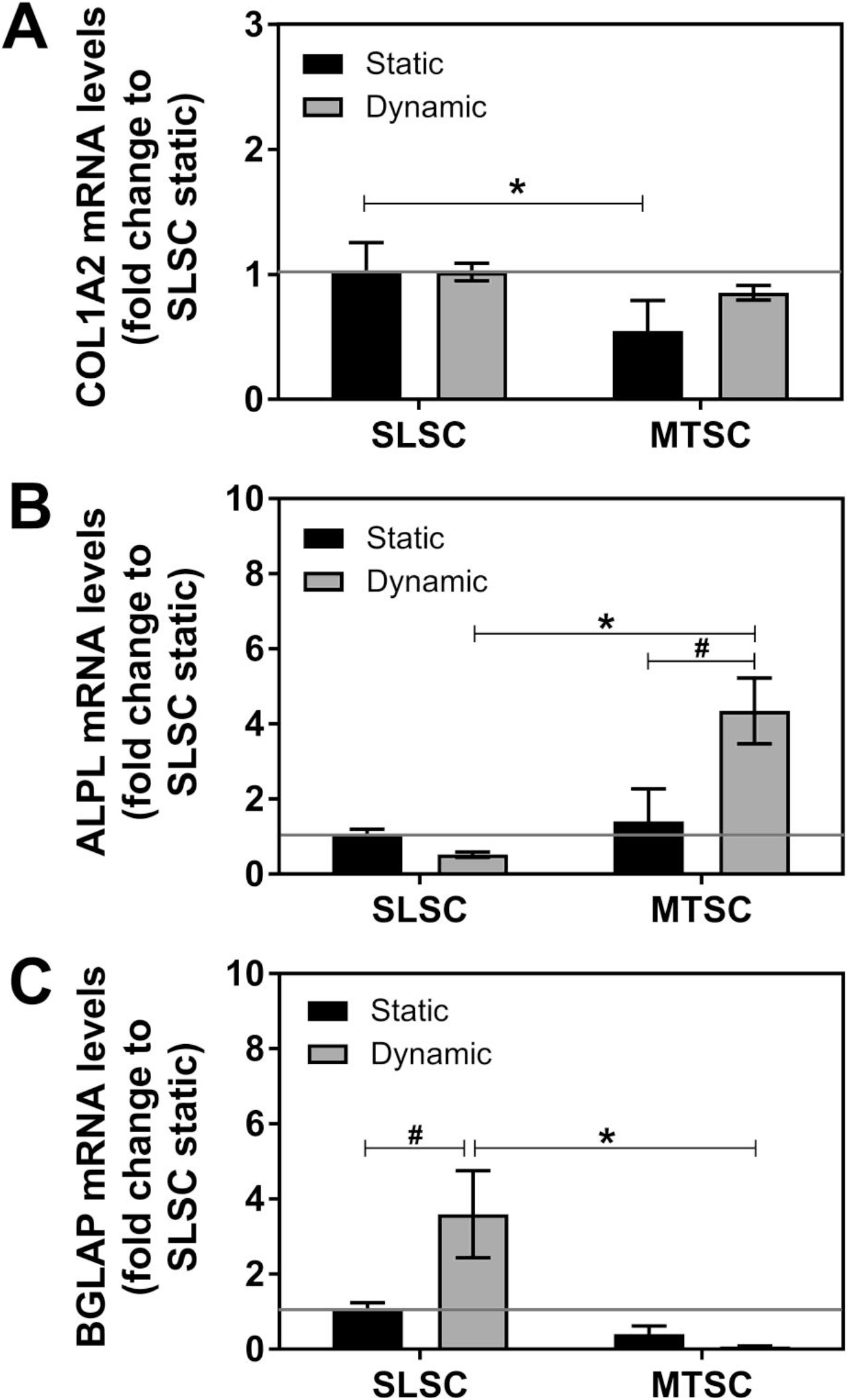
Relative mRNA expression levels of collagen type I (COL1A2) (A), alkaline phosphatase (ALPL) (B), and osteocalcin (BGLAP) (C) in hBMSCs cultured on SLSC or MTSC either under static (black bar) or dynamic (grey bar) culture conditions in osteogenic media for 7 weeks. Values were normalized to housekeeping gene (glyceraldehyde-3-phosphate dehydrogenase [GAPDH]) and expressed as fold change relative SLSC cultured under static conditions, which was set to 1. Values represent the mean ± SD (n = 3). Results were considered statistically significant with p≤ 0.05 between scaffold types (*) and culturing conditions (#), respectively.

### 3.4. ECM mineralization

Micro-CT was used to evaluate the influence of scaffold pore geometry and culture condition on mineralized bone-like tissue formation. Figure 8 shows only the mineralized tissue formed in grey when cells were cultured on SLSC and MTSC scaffolds under static or dynamic culture conditions. Both SLSC and MTSC supported mineralized tissue formation under static and dynamic conditions. Dynamic conditions (figure 8I-P) favored a more homogeneously distributed mineralized ECM deposition in both scaffold types, whereas under static conditions (figure 8A-H) mineralized ECM was limited to the edges of the scaffold or the mid-bottom scaffold’ region in SLSC and MTSC, respectively. The engineered mineralized bone-like tissue architecture was quantified by morphometrical parameters (figure 9). No differences in BV/TV were observed between SLSC and MTSC under static conditions (figure 9A). Nevertheless, the influence of dynamic conditions on BV/TV (figure 9A) and on engineered bone-like tissue distribution along the scaffold (figure 8I-P) was affected by scaffold pore geometry. BV/TV was significantly higher for MTSC under dynamic conditions (12.27% ± 0.58%) compared to SLSC under dynamic conditions (6.17% ± 1.63%, p = 0.004). BS/BV was increased significantly in SLSC under dynamic conditions (24.05 ± 2.29 mm^−1^) compared to SLSC under static conditions (18.46 ± 1.03 mm^−1^, p = 0.016). Lower BS/BV values relate to more compact and plate-like bone structures. In line with the BS/BV data, SLSC under dynamic conditions were characterized by a more homogenous distribution of the mineralized tissue as compared to a compact and dense structure in SLSC under static conditions. Despite the more homogeneous distribution of mineralized tissue and the high BS/BV values (figure 9B), dynamically cultured SLSC scaffolds also showed a dense and compact bone-like surface density at the mid-bottom region of the scaffold (figure 8K). However, this local lower porosity was not as prominent as the overall homogenous distribution of the mineralized tissue throughout this scaffold. No differences were observed for BV/TV and BS/BV values between MTSC under static and dynamic conditions.

**Fig. 8.**
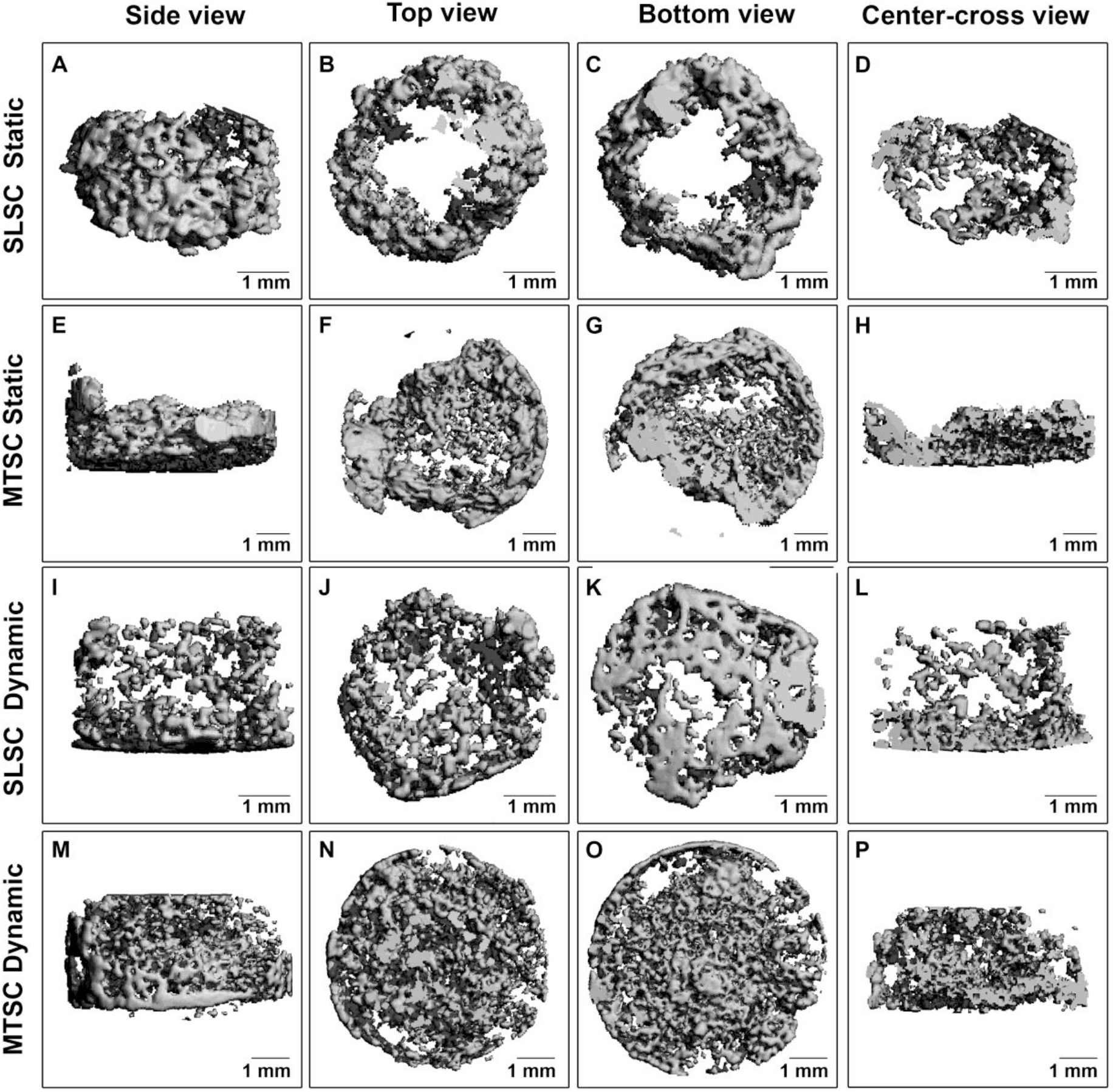
Morphology of engineered bone-like tissue from hBMSCs cultured on SLSC (A-D, I-L) and MTSC (E-H, I-L) under static (A-H) or dynamic (I-P) conditions for 7 weeks. Side view (A, E, I, M) top view (B, F, J, N) bottom view (C, G, K, O) and center-cross view (D, H, L, P). Scale bar lengths are 1 mm.

**Fig. 9.**
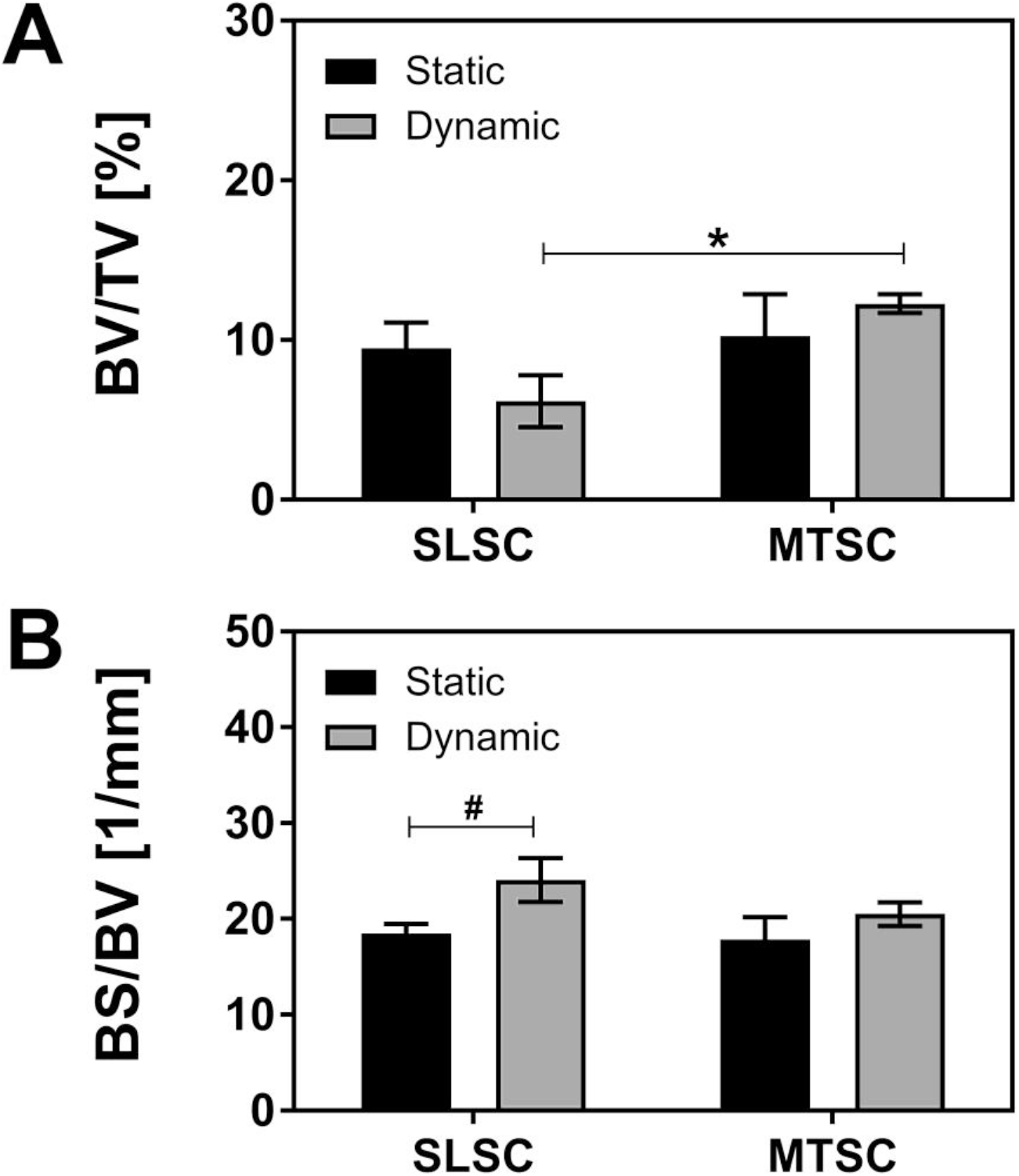
Effect of pore geometry and culture conditions on engineered mineralized bone-like tissue volume fraction (BV/TV) (A) and bone-like tissue surface-to-bone volume (BS/BV) (B). (*) p ≤ 0.05 versus SLSC for each culture condition (#) p ≤ 0.05 versus static condition.

Similarly, the influence of dynamic culture conditions on the trabecular morphometric indices was dependent on scaffold pore geometry (figure 10). In accordance to the BS/BV data, Tb.Th. was significantly smaller in SLSC under dynamic conditions (0.08 ± 0.008 mm) compared to SLSC under static conditions (0.11 ± 0.006 mm, p = 0.054) (figure 10A). Similarly, Tb.Th showed a trend to be smaller in MTSC cultured in dynamic condition compared to MTSC static conditions, although this was not statistically different. Tb.N. (figure 10B) was higher in MTSC under dynamic conditions (1.26 ± 0.07 mm^−1^) compared to MTSC under static conditions (0.9 ± 0.18 mm^−1^, p = 0.087) and significantly higher than in SLSC under dynamic conditions (0.73 ± 0.15 mm^−1^, p = 0.004). Conversely, the Tb.Sp. was lower in MTSC under dynamic conditions compared to MTSC under static conditions and SLSC under dynamic conditions, although differences were not statistically significant (figure 10C).

**Fig. 10.**
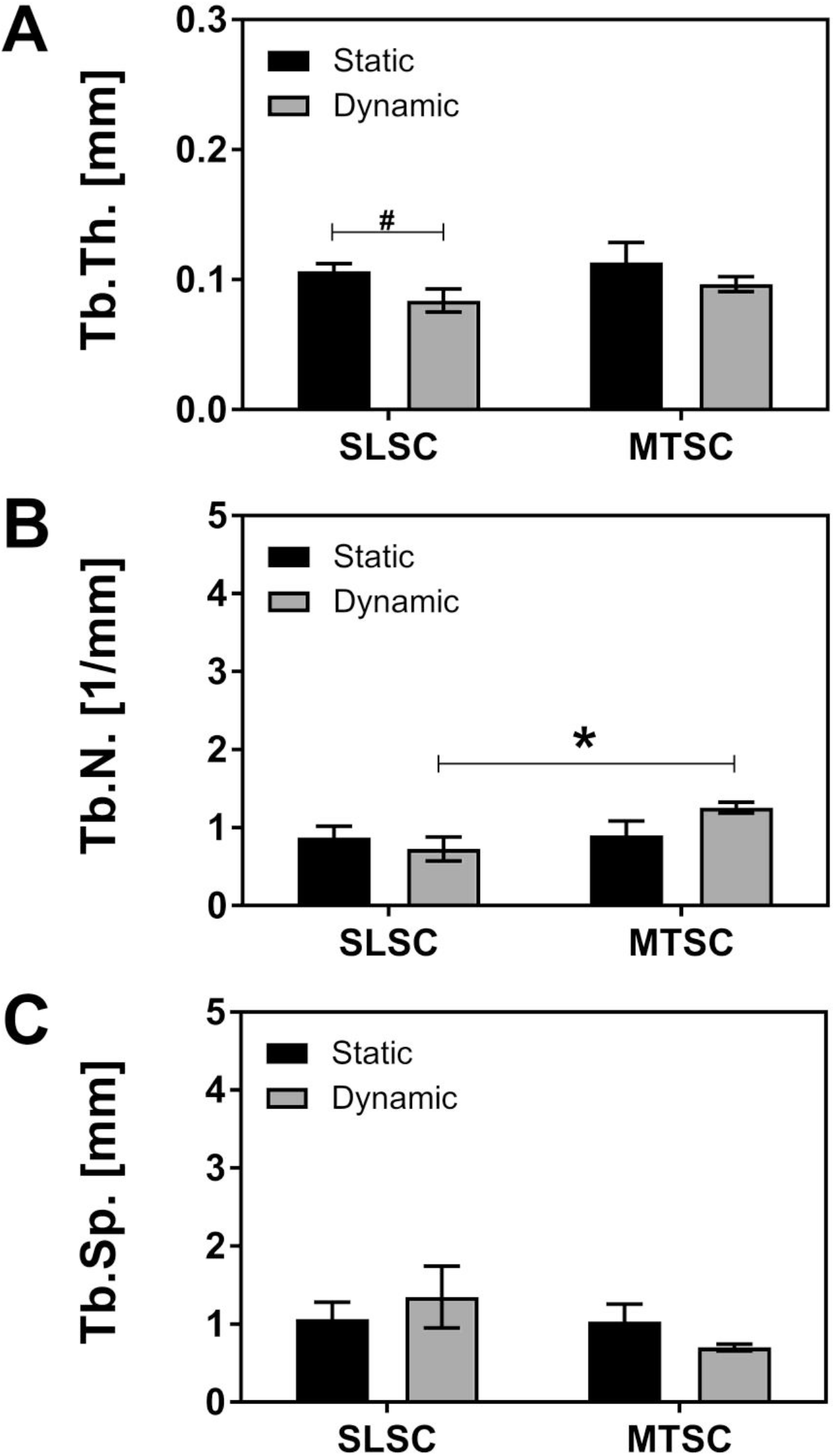
Effect of pore geometry and culture conditions on trabecular structures. Trabecular thickness (Tb. Th) (A), trabecular number (Tb.N) (B) and trabecular separation (Tb.Sp) (C). hBMSCs were cultured on SLSC or MTSC under static or dynamic conditions for 7 weeks. (*) p ≤ 0.05 versus SLSC for each culture condition (#) p ≤ 0.05 versus static condition.

## 4. Discussion

In this study, we used a manufacturing method to systematically fabricate scaffolds with highly controlled monodisperse porous architecture. The aim of the current study was to compare two scaffold manufacturing methods on how scaffold pore geometry influences the response of cells to culture conditions. Evaluated were the distribution and osteogenic cell differentiation as well as the amount, structure and spatial distribution of the mineralized tissue formed. Dynamic culture conditions were used to overcome the nutrient flow limitations and to apply mechanical loading through WSS. Cells (hBMSCs) differentiated towards the osteogenic lineage when cultured either under static or dynamic conditions for both scaffold pore geometries. Nevertheless, distinct responses to fluid flow in gene expression, the amount, structure and spatial distribution of the mineralized ECM formed were shown to be dependent on scaffold pore geometry.

The applied SLSC and MTSC specimens were produced following the same procedure described by Sommer *et al* 2016 [23]. Geometrical features reported by the authors were thus used here to aid the interpretation of the obtained results [23]. Despite their similar local stiffness and a comparable mean pore diameter of about 200 μm, SLSC and MTSC differed in the pore diameter distribution and pore geometry. This resulted in a lower porosity in the MTSC (84.7 ± 2.1 %) compared to SLSC (92.8 ± 0. 6%) (figure S1) [23]. We used histological staining’s to evaluate at the influence of these differences of scaffold porosities on cell proliferation capabilities. The histological results showed that cells could adhere well on both scaffold types but that the ability of cells to proliferate was reduced in the MTSC scaffolds under static conditions, which could indeed be due to a lower pore interconnectivity. Similarly, previous studies have shown that SF scaffolds with similar pore diameter and porosities were able to better support cell proliferation *in vitro* [9, 48, 49]. Assuming a similar MTSC porosity as described by Sommer *et al* [23], the lower porosity and the concave nature of the MTSC might have limited the initial cell distribution throughout the scaffold volume during the cell seeding and persistent growth along the pore surface in MTSC [50]. Nevertheless, this restriction was changed and fluid–induced mechanical stimulation could compensate if the MTSC were cultured under dynamic conditions, as shown from the CFD simulation. Besides enhancing cell distribution, dynamic conditions also significantly increased mRNA levels of ALPL and BGLAP in MTSC and SLSC, respectively, compared to static conditions after 7 weeks of culture. These results indicate that cells cultured under dynamic conditions were at a more mature osteoblastic differentiation stage compared to cells cultured under static conditions. Accordingly, previous reports have shown positive effects of turbulent flow created in spinner flask bioreactors on osteogenic cell differentiation *in vitro* [51, 52]. Although dynamic cultures supported a similar and homogenous distribution of cells for both scaffolds types, COL1A2 gene was stronger expressed in SLSC compared to MTSC under static conditions. The formation of a denser collagenous ECM was also observed in SLSC compared to MTSC under dynamic conditions. In line with this observation, SLSC enhanced osteogenic cell differentiation as shown by an increase in the BGLAP mRNA levels (the hallmark of mature osteoblasts) compared to MTSC under dynamic conditions. Differences in the up-regulation of mid (ALPL) or late (BGLAP) differentiation markers between MTSC and SLSC under dynamic conditions suggest that cells on SLSC were at a more matured differentiation stage. The magnitudes of mechanical forces have been shown to influence gene expression in osteoblastic cells [53]. According to the computational model, the SLSC had a higher surface area fraction (98.21%) exposed to the WSS range under dynamic conditions for osteogenesis than the MTSC (82.3%). In line, dynamic SLSC upregulated the expression BGALP which is a late osteogenic-related marker involved in the bone mineralization. While the underlying mechanisms are not fully clear, we speculate that cells reacted distinctly to the different WSS developed in the two pore geometries.

Osteoblast cell differentiation is characterized by the formation of a mineralized ECM. Although bone-like tissue was formed in all groups, the amount, the structure and distribution of the mineralized ECM was dependent on the pore geometry and whether the constructs were grown statically or dynamically. Interestingly and in contrast to the gene expression profile observed, MTSC under dynamic conditions had a significantly increased BV/TV compared to SLSC under dynamic conditions. This is in agreement with our observation that pore geometry can lead to differences in the bone-like structure along the scaffold under dynamic conditions. On the other hand, the computational prediction showed that a larger surface area fraction of the SLSC was exposed to a WSS range for stimulating mineralization than that of the MTSC, which is in contrast to results obtained experimentally. A major limitation of this study is that the CFD model was based on idealized geometrical scaffold features created by CAD, assuming an empty scaffold where tissue growth fills the pores overtime. Specimen specific CFD models created directly from micro-CT images may be a solution to assess local WSS in the scaffold micro-environment. Nevertheless, such models are currently still hampered by the fact that the micro-CT images only show mineralized tissue and remnants of the scaffolds and newly formed soft tissue filling in the pores cannot be identified precluding identification of the “true” pore size and the associated fluid flow through the pore network using the CFD model. A further limitation of the current approach using idealized CFD models, is that scaffolds used experimentally display irregular pore shapes causing variations in pore size, which results in WSS discrepancies between real and idealized scaffold pore geometries. A previous CFD study compared WSS within (i) irregular SF scaffolds with a porosity of 55% and an average pore size of 160 μm and (ii) regular PCL scaffold with a porosity of 38% and an average pore size of 220 μm [54]. It was found that the resultant average WSS difference was about 19% between these two scaffolds [54]. In this study, the CAD scaffold geometry has the same average pore size and porosity as the real scaffold, therefore, it is likely that the WSS difference will be smaller (e.g. < 19%). Furthermore, by using an idealized scaffold geometry for the CFD model, it is expected that the CFD results are not only supporting this study, but also informative for other tissue engineering studies, which will use scaffolds made by different methods (e.g. 3D printing) but with similar porous geometric features in similar environments. Nevertheless, CFD results based on empty scaffolds will be applicable only for the early phase of the experiment (e.g. day 0). Recent computational studies have shown that cell/tissue growth within a scaffold can change the mechanical stimulation (e.g. WSS) on cells [55]. Moreover, the tissue morphologies had a distinct influence on the resultant WSS on cells [55]. Such change of WSS would further influence the ECM formation and mineralization. Our histological results have revealed that in MTSC only a few cells were laying on the scaffold surface, which is the place where the mechanical stimulation is expected to happen in the computational model. Based on the 3D micro-computed tomography images, MTSC under dynamic conditions favored a more homogeneously distributed ECM mineralization characterized by a more trabecular bone-like structure including locally restricted areas of plate-like bone (larger surface of compact bone), compared to the rest of the groups. Compared to the spherical nature of PCL particles used for making the pores in MTSC, the rock-salt crystal structure of NaCl rendered more faceted pore surfaces in the SLSC scaffolds. It has been reported that surface concavities act as a strong template for *in vitro* surface mineralization, even in the absence of living cells [28]. Surface mineralization is probably favored within pore concavities as compared to planar surfaces because nuclei are less exposed to shear and are thus able to grow larger in sites where the local fluid velocity is low [19, 28]. We hypothesize that MTSC with more regular and concave surfaces favor a more homogeneous deposition of mineral due to inhibited fluid motion close to regularly distributed pore walls. By contrast, the more planar features present in SLSC might result in an unevenly distributed mineral deposition with higher mineral formation at concave surfaces and lower mineral formation at planar surfaces of the pore. Differences in fluid flow next to concave and planar surfaces might then modulate the local ion concentration affecting the nucleation process and therefore mineral tissue formation. The decrease in BV/TV for SLSC under dynamic compared to static conditions could be due to the higher flow velocities at planar surfaces, which in turn could lead to more fluctuations in the local concentrations of Ca^2+^ and PO_4_^3^, as previously suggested by Bianchi *et al* 2014 [28]. This effect might be enhanced by the high porosity of the SLSC [23].

In conclusion, our results suggest that scaffold pore geometry and porosity generate different flow velocities/WSS within the scaffolds in the bioreactor, which in turn directly affect cell differentiation and bone-like tissue formation. While SLSC enhanced osteoblast cell differentiation, the spherical and more regular concave pore geometry in MTSC could probably favor nucleation of ions increasing the amount of bone-like tissue formation. These findings point out the role of pore geometry on the stimulation of cell differentiation and indirectly on guiding the mineral deposition process. This suggests the importance of 1) the fluid flow imposed by the scaffold’s pore geometry on the cell response and the formation of an engineered bone-like tissue, 2) control and tune the scaffold pore geometry to modulate the amount and structure of engineered bone-like tissue. Fabrication of complex scaffolds, by additive manufacturing methods, comprising regions with more concave pores with others with more planar pore geometries might thus represent a promising direction towards the tunability of the BTF and to mimic the structural complexity of the bone tissue and its spatial distribution to each specific need. Our findings also demonstrate the relevance of the chosen mechanical loading as loads applied can overrule or affect the influence of the scaffold pore geometry. These conditions should be carefully chosen and included for quality assurance during assessment of new scaffolds. Future studies could further benefit from the inclusion of specimen specific CFD models, which would allow a more direct correlation of fluid dynamics, complex scaffold architectures and tissue response as assessed from time-lapsed high-resolution micro-CT scans of the mineralizing scaffolds.

## 5. Acknowledgements

The authors thanks to Trudel Inc. for the supply of the silk cocoons and ScopeM for providing access to their histology equipment. The research leading to these results has received funding from the European Union Seventh Framework Programme (FP7/2007-2013) under grant agreement n°. 329389, 262948 and 336043.

## Supplementary material

**Fig. S1.**
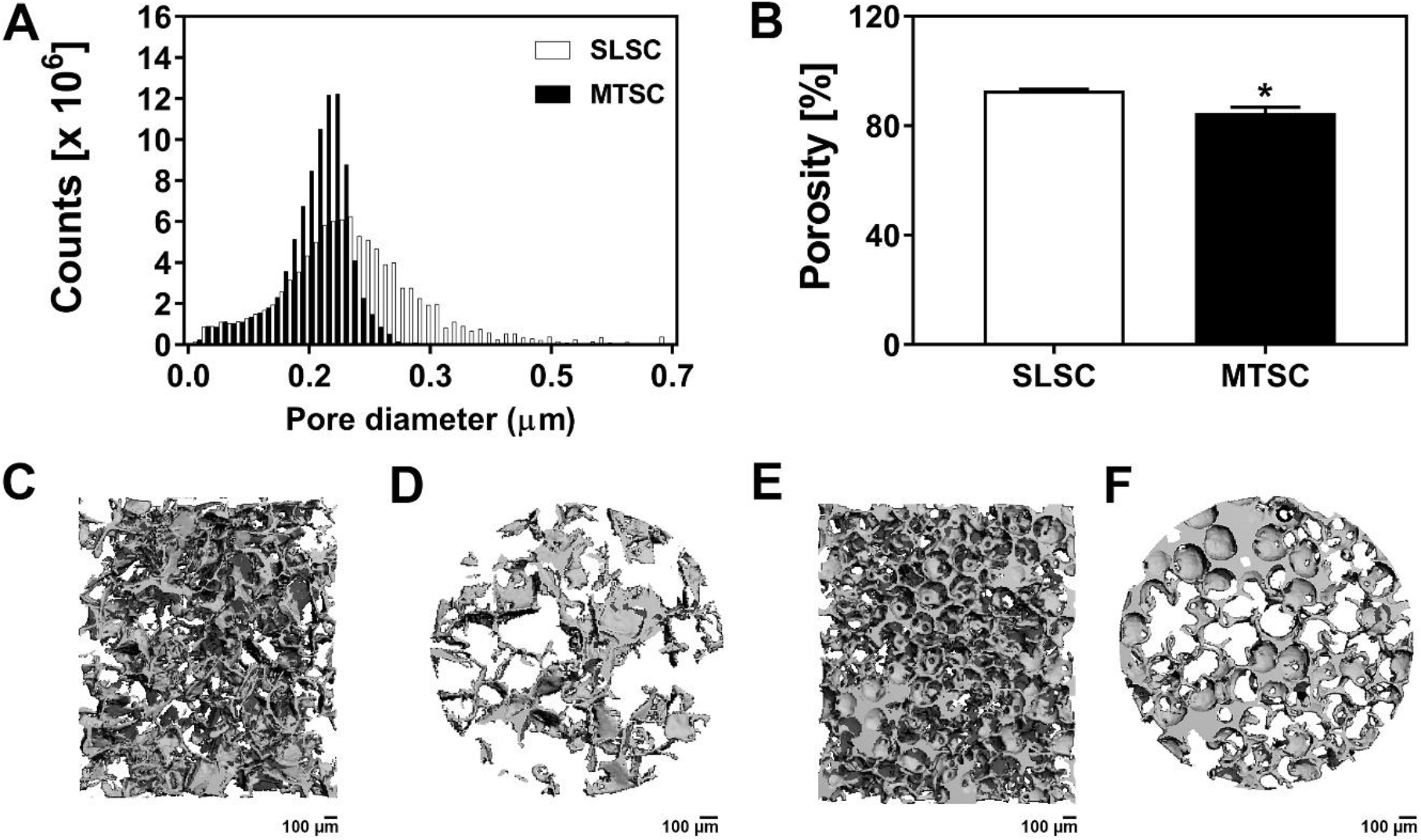
Geometrical features of SLSC and IOSC. Quantitative pore diameter distribution (A) and porosity (B). Micro-CT images from a vertical (C, E) and horizontal (D, F) cut-plane of SLSC (C, D) and IOSC (E, F). Scale bar lengths are 100 μm. Adapted from Marianne Sommer *et al.* (2016) [23].

**Fig. S2.**
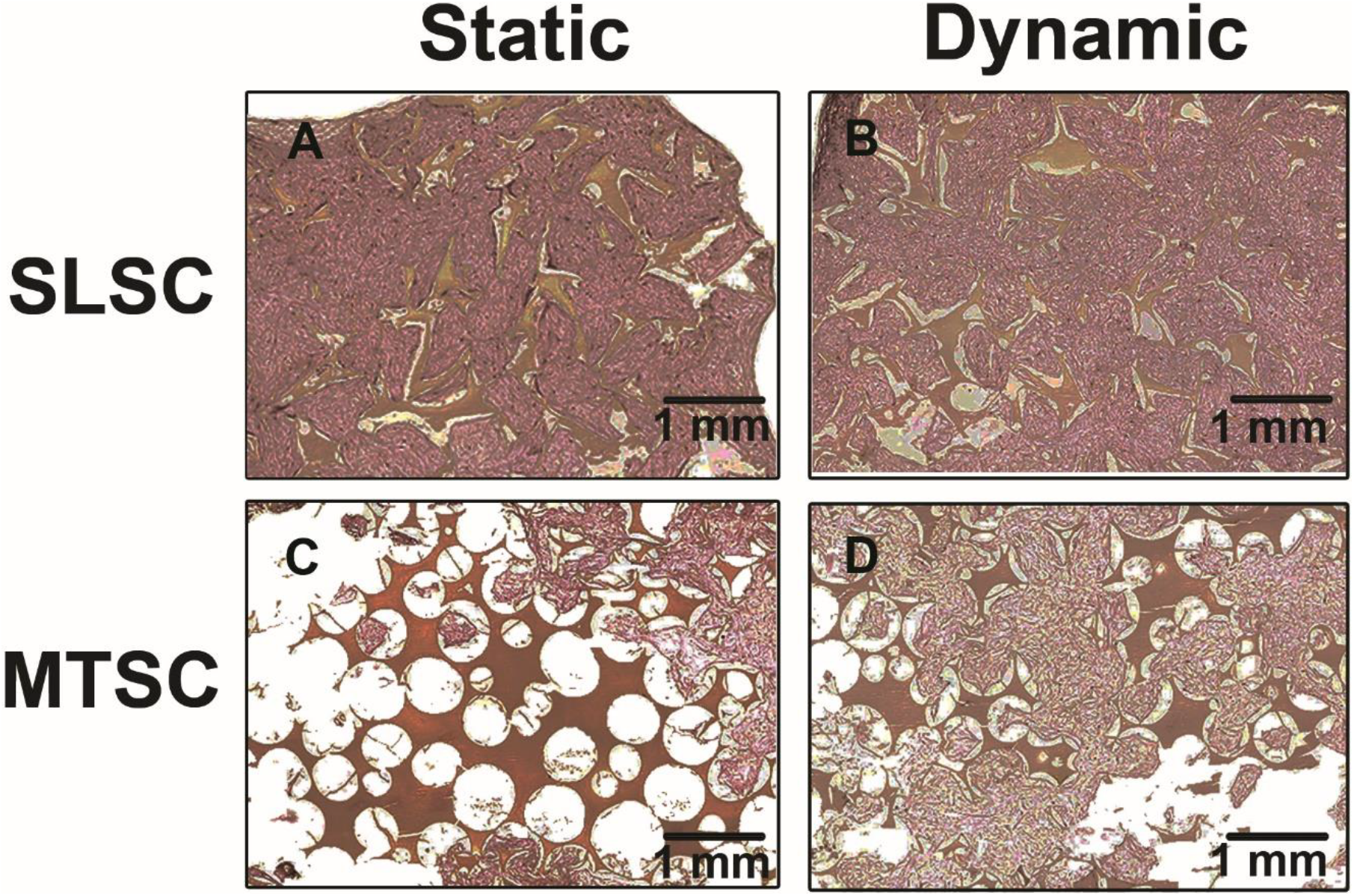
Magnified view of the formation of a collagenous ECM in the middle region of the scaffolds. Combined Weigert’s haematoxylin staining (cell nuclei - dark purple spots) and Sirius red staining (collagenous ECM - pink) on hBMSCs cultured on SLSC (A, B) and MTSC (C, D) either under static (A, C) or dynamic conditions (B, D). Scaffolds are shown in red color. Scale bar lengths are 1 mm.

